# Impact of cross-validation designs on cattle behavior prediction using machine learning and deep learning models with tri-axial accelerometer data

**DOI:** 10.1101/2025.01.22.634181

**Authors:** Jin Wang, Ziwen Yu, Ricardo C. Chebel, Haipeng Yu

## Abstract

Assessing cattle behaviors provides insights into animal health, welfare, and productivity to support on-farm management decisions. Wearable accelerometers offer an alternative approach to traditional human evaluation, providing a more objective and efficient method for predicting cattle behavior. Random cross-validation (CV) is commonly used to evaluate behavior prediction by splitting data into training and testing sets, but it can yield inflated results when records from the same animal are included in both sets. Block CV splits data by block effects, offering a more realistic evaluation but remains underexplored for predicting multi-class imbalanced cattle behavior. Additionally, deep learning (DL) models have not been fully explored for behavior prediction compared to machine learning (ML) models. The objectives of this study were to examine the impact of CV designs on multi-class imbalanced cattle behavior prediction and to compare the performance of ML and DL models. Three ML and two DL models were used to predict the four behaviors of six beef cows from a public tri-axial accelerometer dataset, with model performance evaluated using both hold-out and leave-cow-out CV designs representing random and block CV, respectively. In hold-out CV, ML models achieved accuracies of 0.94 to 0.96 and F1 scores of 0.93 to 0.95, while DL models achieved accuracies of 0.9 to 0.92 and F1 scores of 0.89 to 0.91. In the leave-cow-out CV, ML models obtained accuracies of 0.72 to 0.82 and F1 scores of 0.64 to 0.82, whereas DL models obtained accuracies of 0.76 to 0.82 and F1 scores of 0.64 to 0.76. Generally, ML models outperformed DL models in the hold-out CV, but the multi-layer perceptron DL model demonstrated comparable or superior performance in the leave-cow-out CV. All models performed better with hold-out CV than leave-cow-out CV. Our results suggest that CV designs can affect behavior prediction performance. While a random CV produces seemingly good predictions, these results can be artificially inflated by the data partition. A block CV that strategically partitions data could be a more appropriate design.

## 2 Introduction

Behavior pattern changes in dairy cattle have been reported to be associated with respiratory disease [1] and lameness [2], estrus [3], and welfare[4]. Monitoring behavior changes can provide insights into improving cattle health and general well-being. Traditional on-farm monitoring approaches have relied on human observations, which are labor- and time-intensive and prone to subjective errors [5]. To address these limitations, various precision livestock farming technologies have been developed to facilitate objective, non-intrusive, and real-time cattle behavior monitoring [6, 7, 8, 9]. Among these technologies, wearable accelerometer sensors are most commonly used for monitoring and predicting the behavior of beef and dairy cattle [10, 11, 12] and other species [13, 14]. Typically, the generated raw accelerometer data are first preprocessed and then fitted into various prediction models for behavior predictions. To evaluate the performance of behavior prediction based on accelerometer data, previous studies have commonly applied cross-validation (CV) to partition the dataset into training and testing sets. The training set is used to fit the model, while the testing set is used to assess the prediction performance of the fitted model. Most of these studies have applied random CV, such as hold-out or k-fold CV, which randomly splits the whole dataset into training and testing sets to evaluate prediction performance [e.g., 15, 16]. However, accelerometer data are typically recorded sequentially over time for each cattle, but random CV does not account for this inherent structure when splitting data, potentially assigning data from the same cattle to both the training and testing sets. This can introduce dependencies or data leakage between the sets due to the covariance introduced by the same cattle, potentially resulting in overly optimistic and inflated prediction performance due to overfitting [17, 18]. An alternative approach is block CV, which strategically partitions data by factors such as animal or farm to account for the inherent data structure and has been applied in other fields. For example, Wang and Bovenhuis [18] examined the use of milk infrared spectra to predict methane emissions using both random and block CV and reported that random CV produced overly optimistic predictions compared to block CV, likely due to farm-specific confounding effects. More recently, a study applied accelerometer data to predict grazing versus non-grazing behaviors using both random and block CV and highlighted the importance of using management knowledge to strategically split data [19]. However, studies on the impact of CV designs on multi-class imbalanced behavior prediction in beef cattle remain scarce in the literature.

Although machine learning (ML) models have been extensively applied to predict cattle behavior from accelerometer data, deep learning (DL) models such as a convolutional neural network (CNN), which is commonly utilized in computer vision for livestock farming systems [20, 21, 22], remain relatively unexplored. To the best of our knowledge, previous work has only compared ML and DL models in predicting cattle behavior using random CV [23, 24], and no study has compared their prediction performances using more realistic block CV.

In this study, we examined how different CV designs affect the results of ML and DL-based models to predict cattle behavior from accelerometer data. Specifically, our objectives were to 1) assess the impact of CV designs on the prediction of multi-class imbalanced cattle behavior, and 2) compare the prediction performance of ML and DL models using both random and block CV.

## 3 Materials and Methods

### 3.1 Tri-axial accelerometer sensor data

The tri-axial sensor data utilized in this study were sourced from a public dataset [25]. These data were collected from six beef cows via a 16-bit *±* 2*g* Kionix KX122-1037 accelerometer attached to their necks at a sampling frequency of 25 Hz. Meanwhile, video data were recorded and annotated by humans by aligning the timestamps of the accelerometer data with the video recordings. This process resulted in annotations for 13 distinct behaviors. After removing missing records, a total of 22, 987 records across 197 min for six beef cows were retained for analysis in this study. Four major behaviors used in this study were resting while standing (RES), ruminating while standing (RUS), moving (MOV), and grazing (GRZ). The number of records for each behavior is shown in Figure 1.

**Figure 1:**
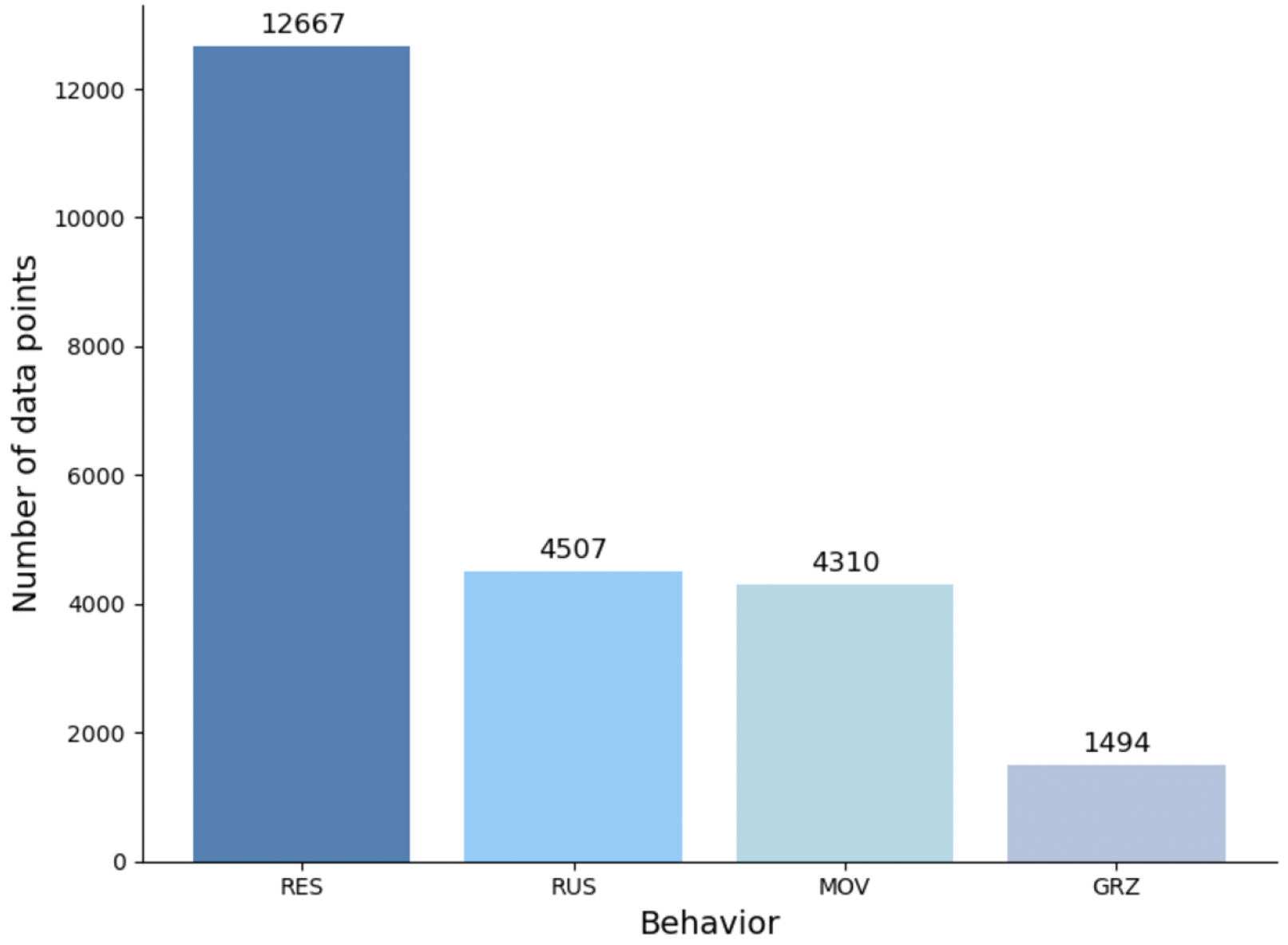
Histogram showing the number of data points for each of the four cattle behaviors. RES, RUS, MOV, and GRZ refer to resting while standing, ruminating while standing, moving, and grazing, respectively.

### 3.2 Data preprocessing and feature extraction

The accelerometers captured three raw features that measured the instantaneous acceleration of cattle along the axes of X, Y, and Z. From these raw features, we derived four additional instantaneous features, including the magnitude of acceleration, pitch angle, roll angle, and signal magnitude area. These four features were calculated using the formulas detailed in Table 1. To improve data quality and computational efficiency, we applied data downsampling to the seven instantaneous features using a sliding window method. A window size of 1 second with a step size of 0.5 seconds (Figure 2) was used to compute summary statistics, including the mean, minimum, maximum, and variance for each instantaneous feature. This process resulted in a total of 28 features. We further excluded 8 features showing a high correlation coefficient (≥ 0.9) with at least two other features to address the collinearity among features. The remaining 20 features, as shown in Table 1, were used to develop models for predicting four cattle behaviors.

**Table 1:**
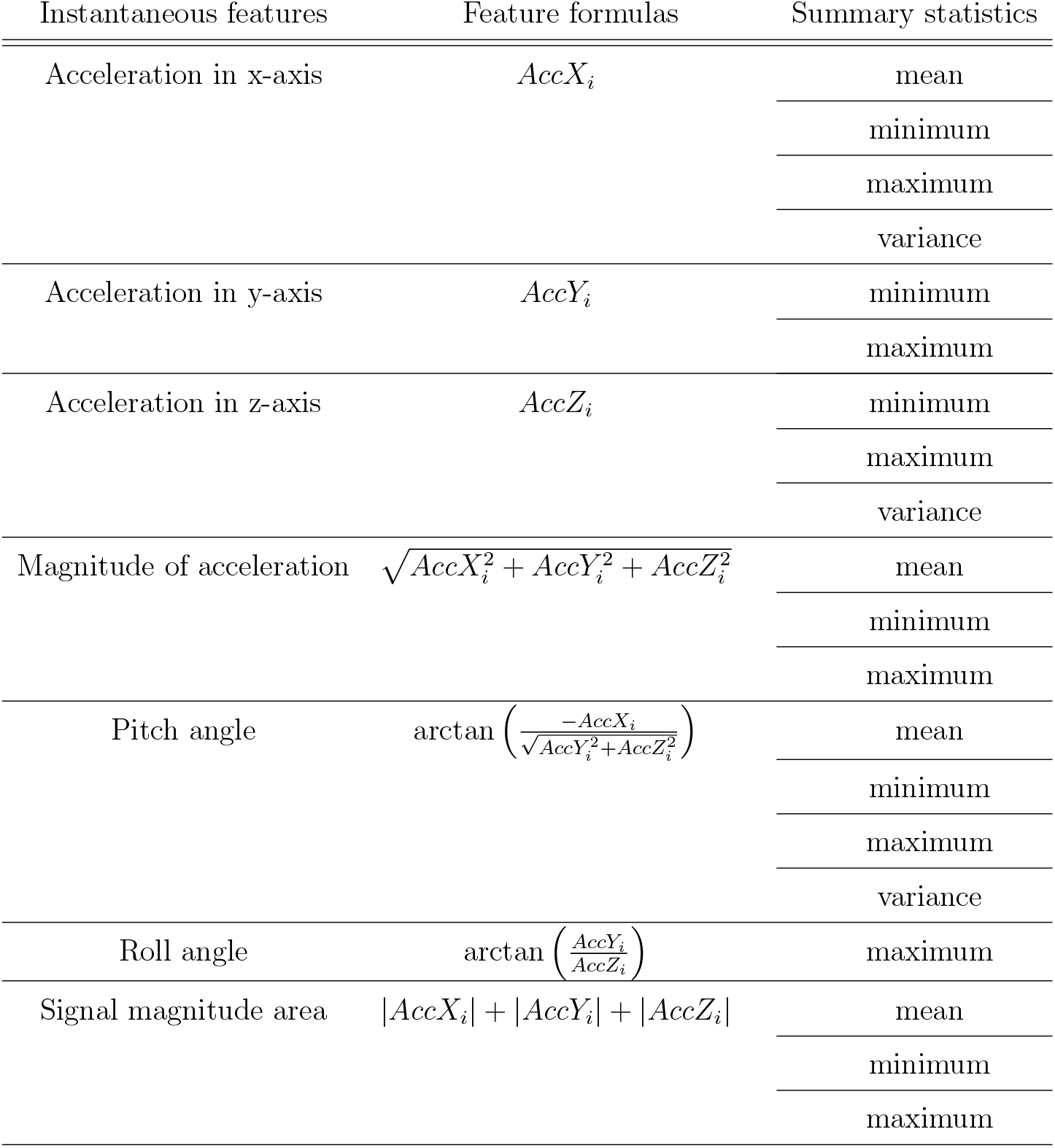
Seven instantaneous features with their corresponding formulas and 20 summary statistics extracted from these features. Here, *i* represents the instantaneous feature measured at the *i*-th time point.

**Figure 2:**
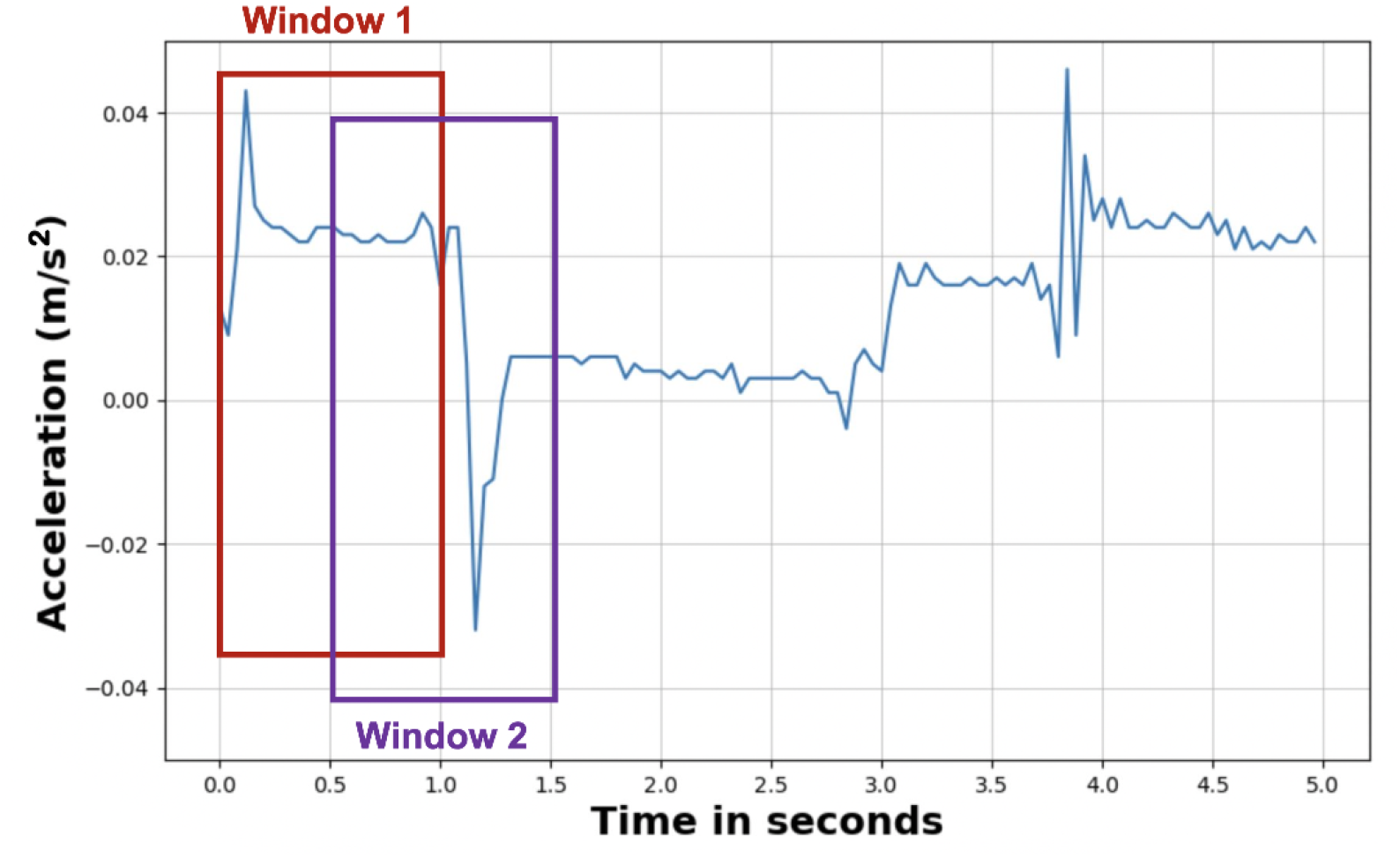
Illustration of the sliding window method applied to acceleration data, with a window size of 1 second and a step size of 0.5 seconds.

### 3.3 Cattle behavior prediction models

#### 3.3.1 Machine learning models

Three ML models including k-nearest neighbors (KNN), random forest (RF), and gradient boosting (GB) were used to predict four cattle behaviors. The KNN is a non-parametric supervised ML model that classifies an input data point based on its distance from the nearest k neighbors [26, 27]. The final class of the input data, representing the type of cattle behavior in this case, is determined through majority voting over the behaviors of these k neighbors. The RF and GB are both supervised ensemble learning methods used for classification. The RF employs parallelized training by creating multiple decision trees from random subsets of data and features, with the final class determined by the mode of the results from the individual trees[28]. In contrast, the GB builds ensemble trees sequentially, where each new tree learns from and corrects the gradients (errors) of the predictions made by the preceding tree in the series [29]. The hyperparameters of all three ML models were tuned using CV as described in the following section to improve model performance. The details of the best hyperparameters selected through CV are provided in Table 2. All ML models were implemented using the scikit-learn Python package [30].

**Table 2:** Summary of hyperparameter tuning results for machine learning models and multilayer perceptron model.

| Model hyperparameters | HO 5:5 | HO 6:4 | HO 7:3 | HO 8:2 | HO 9:1 | L1CO | L2CO | L3CO |
| --- | --- | --- | --- | --- | --- | --- | --- | --- |
| *KNN* |  |  |  |  |  |  |  |  |
| number of neighbors | 3 | 4 | 4 | 4 | 4 | 22 | 46 | 99 |
| *RF* |  |  |  |  |  |  |  |  |
| max depth | 20 | 40 | 30 | 30 | 30 | 10 | 40 | 10 |
| min samples for splitting | 5 | 5 | 5 | 5 | 5 | 15 | 15 | 10 |
| number of trees | 200 | 200 | 300 | 300 | 300 | 300 | 200 | 250 |
| *GB* |  |  |  |  |  |  |  |  |
| contribution of each tree | 0.5 | 0.5 | 0.5 | 0.5 | 0.5 | 0.1 | 0.1 | 0.01 |
| max depth | 8 | 8 | 8 | 8 | 8 | 8 | 8 | 5 |
| min samples per leaf | 3 | 3 | 1 | 1 | 1 | 1 | 2 | 1 |
| min samples for splitting | 2 | 2 | 4 | 6 | 4 | 2 | 6 | 2 |
| number of boosting stages | 150 | 150 | 150 | 150 | 150 | 150 | 150 | 100 |
| *MLP* |  |  |  |  |  |  |  |  |
| number of layers | 3 | 3 | 2 | 3 | 3 | 2 | 3 | 3 |
| learning rate | 0.0005 | 0.0005 | 0.0005 | 0.001 | 0.001 | 0.001 | 0.0005 | 0.0005 |
| number of dense unit | 128/128/256 | 128/256/256 | 128/256 | 256/256/128 | 256/256/128 | 256/256 | 128/64/128 | 128/128/128 |
| dropout layers (rate) | F/F/F | F/F/F | F/F | F/F/F | F/F/F | F/T (0.3) | F/F/F | F/F/F |

HO, hold-out cross-validation; L1CO, leave-one-cow-out cross-validation; L2CO, leave-two-cow-out cross-validation; L3CO, leave-three-cow-out cross-validation; KNN, k-nearest neighbor; RF, random forest; GB, gradient boosting classifier; MLP, multi-layer perceptron. The numbers after HO indicate the ratios of training and test sets. F and T refer to False and True, indicating whether the dropout layer is included.

#### 3.3.2 Deep learning models

In this study, we implemented two DL models for predicting cattle behavior. The first is a multi-layer perceptron (MLP), which is also known as a feedforward artificial neural network or a deep feedforward network. The MLP consists of a series of fully connected layers, including input, hidden, and output layers. The relationships between these fully connected layers can be modeled as

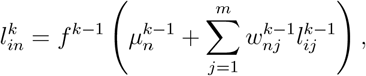

where 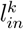 refers to the *n*-th neuron in the *k*-th layer for the *i*-th individual’s record, 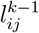 is the *j*-th neuron in the layer 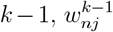 is the weight relating 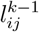 to 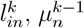 is the intercept (commonly referred to as a “bias” in DL), and *f*^*k*−1^ denotes an activation function that links these two layers. Here, we used a non-linear rectified linear unit (ReLU) as the activation function for linking all layers, except for the output layer, where a softmax function was applied. To optimize the model’s performance, we applied a grid search using CV, as detailed in the following section, to tune the hyperparameters. These hyperparameters included the number of hidden layers, the number of neurons in the hidden layers, the dropout rate, and the learning rate. A categorical cross-entropy loss function and the Adam optimizer were used for model fitting. The details of the tuned hyperparameters in the optimized MLP model are described in Table 2.

The second DL model used in this study is a CNN, which is designed to process more complex data efficiently and accurately by significantly reducing the number of parameters in the model compared to an MLP. A CNN consists of a “backbone” and a “head”, where the former acts as a feature extractor and the latter acts as a classifier. In the feature extractor, a convolution operation is performed using filters that slide across the input data (i.e., 20 sensor features in this study) to extract patterns or information. In detail, each filter has one or multiple kernels, each with a set of weights determined by the window size of the kernel. These weights are used to calculate an output by summing the product of the weights and the input variables within the kernel’s window. An activation function is then applied to the output to obtain activation maps. In this study, we used ReLU as the activation function, except for the last layer, which used a softmax function. To refine the features in these activation maps and reduce their dimensionality, a pooling layer was added after the convolutional layer, taking the maximum values using a sliding filter. The classifier is a fully connected MLP, as described above, which transforms the features extracted by the feature extractor into class probabilities. We tuned the hyperparameters of the CNN using CV (as described below) to optimize the model prediction performance, including the number of convolutional layers, the number of filters and kernel size in each convolutional layer, the number of neurons in the hidden layer of the classifier, and learning rate. The optimized CNN structures are displayed in Figure 3 and Supplementary Figures S1, S2, and S3. All MLP and CNN models were implemented using the open-source ML platform TensorFlow [31].

**Figure 3:**
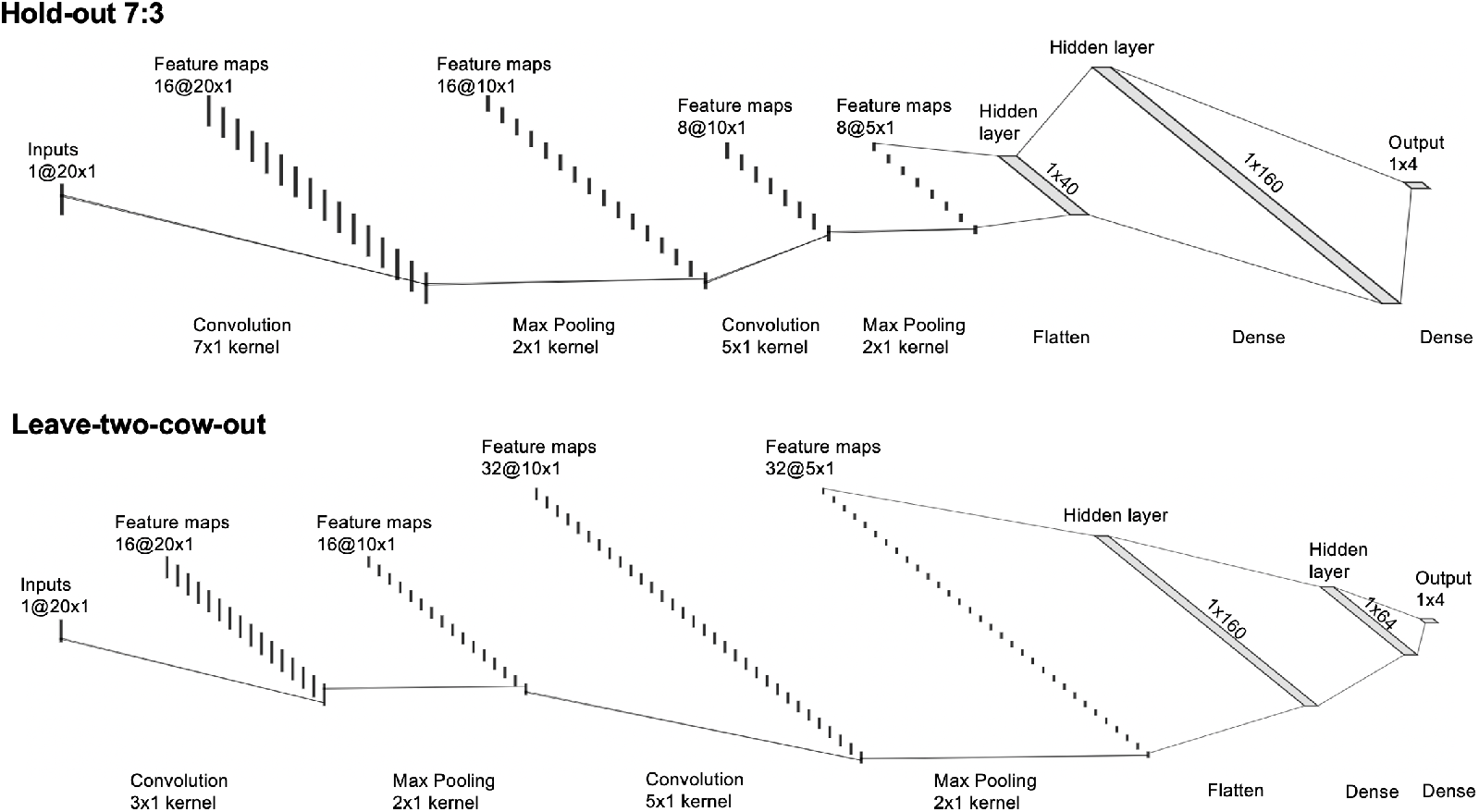
Convolutional neural network structures for hold-out (7 : 3) and leave-two-cow-out cross-validation designs after hyperparameter tuning.

### 3.4 Evaluation of behavior prediction performance

#### 3.4.1 Cross-validation designs

The whole sensor dataset was first partitioned into training and testing sets. The training set was further used to tune the hyperparameters in ML and DL models as described above using a 5-fold CV. Once the hyperparameters were tuned, the parameters of the ML and DL models were estimated using the whole training set and then used to predict the cattle behavior in the testing set. The model performance was evaluated by comparing the predicted behavior with the observed behavior in the testing set using the evaluation metrics described in the following section. In this study, we applied two CV designs to create the training and testing sets. The first is a hold-out CV, a form of a random CV, where the sensor data from six cows for 197 min were randomly partitioned into training and testing sets using five ratios (i.e., 9 : 1, 8 : 2, 7 : 3, 6 : 4, and 5 : 5) as shown in Figure 4. The second CV design applied in this study is a leave-cow-out (LCO) CV, which is a form of block CV. Figure 5 shows an example of leave-one-cow-out (L1CO) CV where in each iteration the sensor data of five cows for 197 min were used to train a model to predict the behavior of the remaining one cow across 197 min. By sequentially excluding one cow as the testing set in each iteration, this process was repeated six times, and the final model performance was the average performance across six iterations. Following a similar procedure, we also applied leave-two-cow-out (L2CO) CV and leave-three-cow-out (L3CO) CV.

**Figure 4:**
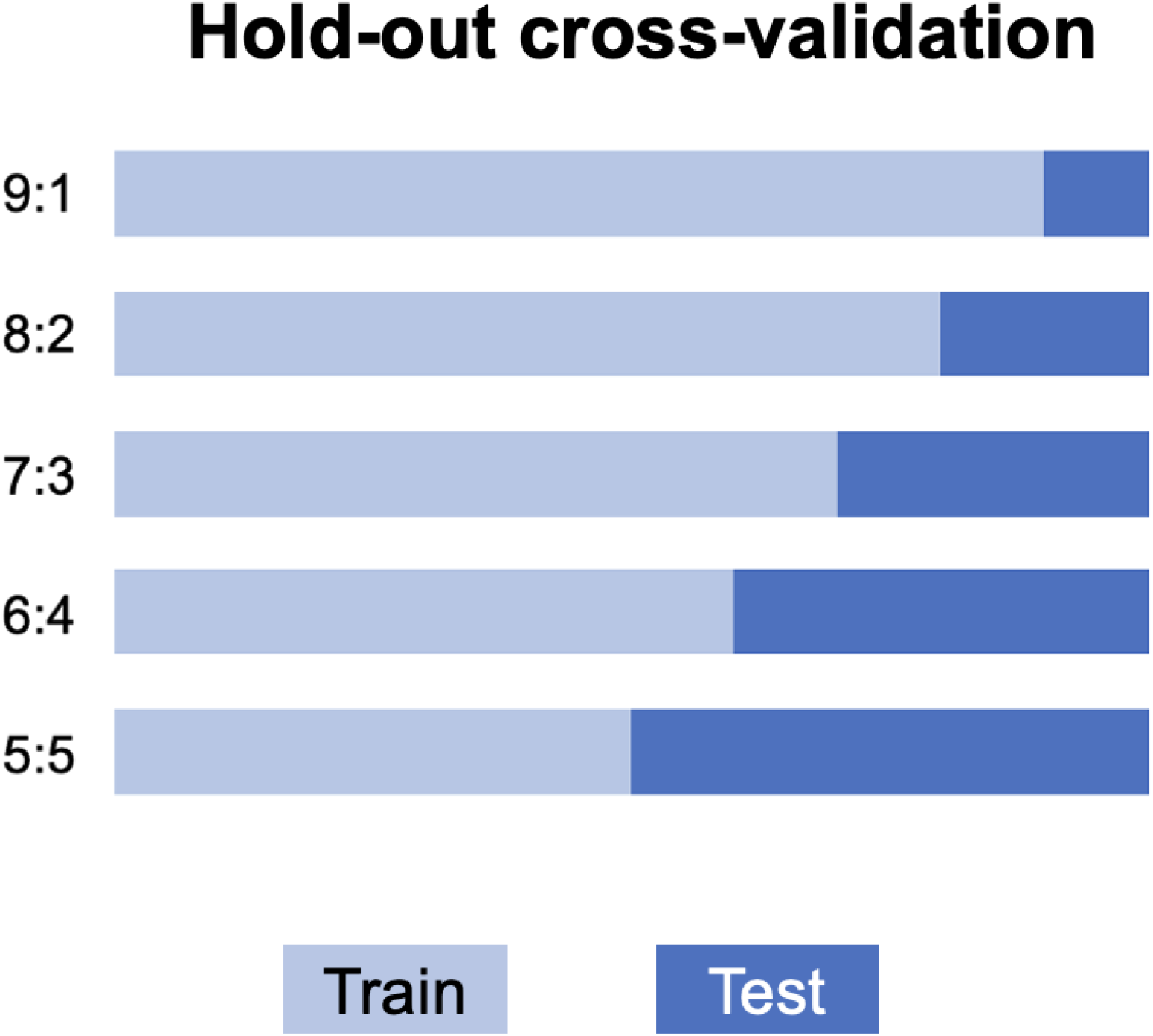
Visualization of five hold-out cross-validation scenarios with varying training to testing ratios including 9 to 1, 8 to 2, 7 to 3, 6 to 4, and 5 to 5. The light blue segments represent the training set proportions while the dark blue segments represent the testing set proportions.

**Figure 5:**
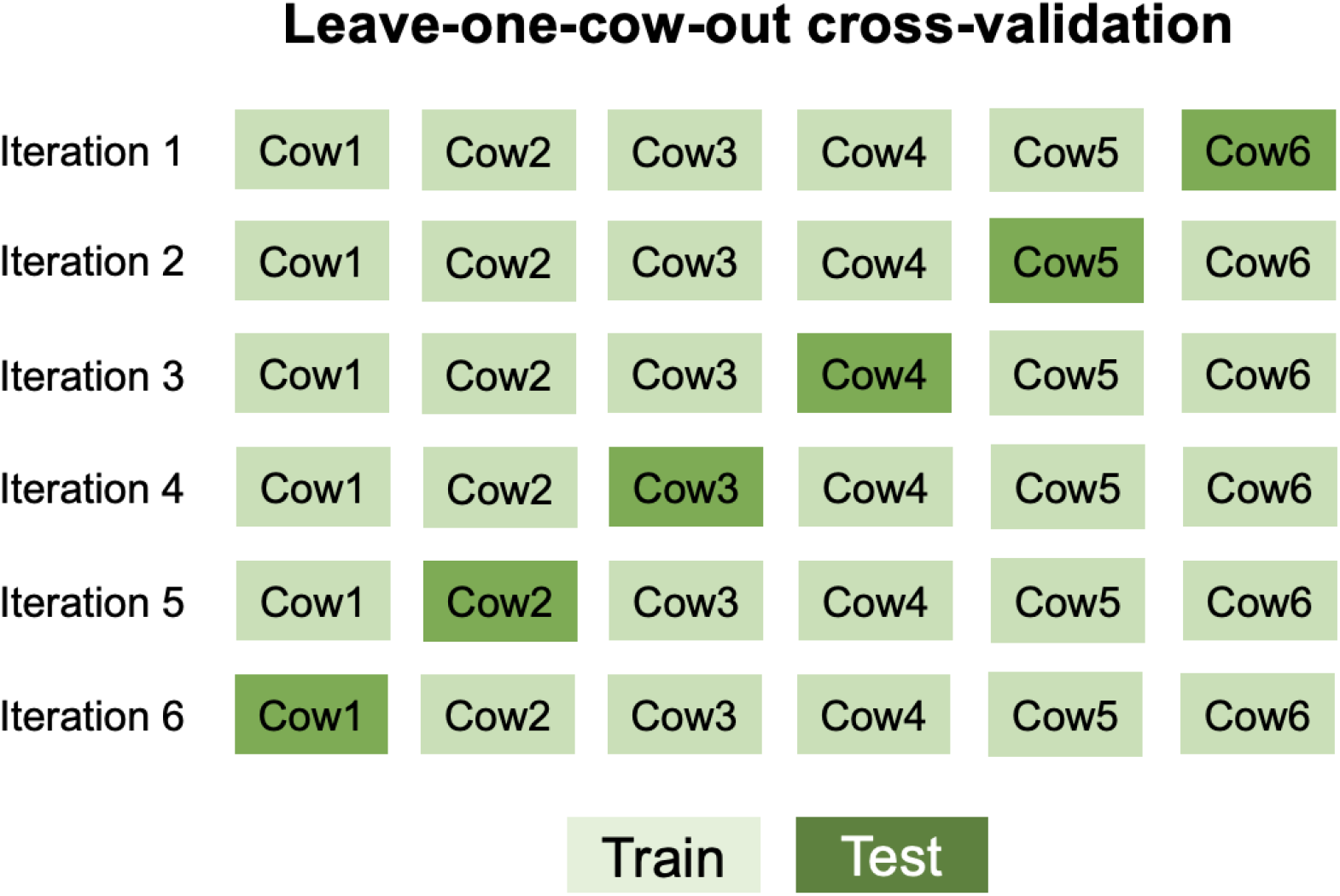
Illustration of the leave-one-cow-out cross-validation design. In each iteration, data from one cow (highlighted in dark green) is assigned to the testing set, while data from the remaining cows (light green) are used for training. This process is repeated until every cow has been used as the testing set once.

#### 3.4.2 Model evaluation metrics

The performance of behavior prediction using ML and DL models was evaluated using four metrics, including accuracy, macro precision, macro recall, and macro F1 score, as detailed below.

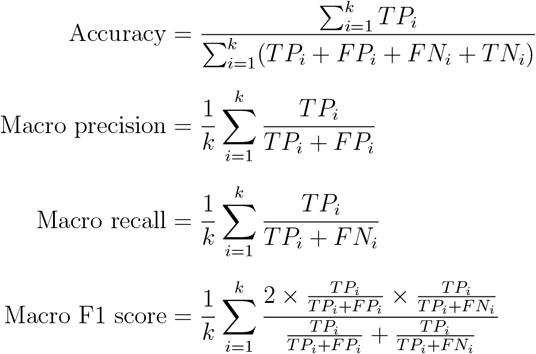

Here, *i* refers to the *i*-th behavior, which ranges from 1 to *k* = 4. *TP*_*i*_ (true positive) is the number of instances correctly classified as behavior *i, FP*_*i*_ (false positive) is the number of instances incorrectly classified as behavior *i* but actually belonging to other behaviors in the ground truth video, *FN*_*i*_ (false negative) is the number of instances belonging to behavior *i* in ground truth video but incorrectly classified as other behaviors, and *TN*_*i*_ (true negative) is the number of instances correctly classified as not belonging to behavior *i*. In this study, we used macro metrics to evaluate the performance of multi-behavior prediction because the cattle sensor data we used are imbalanced (Figure 1), and we are interested in the overall prediction performance of all behaviors.

## 4 Results

The performance of the ML and DL models across two CV designs for the prediction of beef cattle behavior was evaluated using accuracy, macro precision, macro recall, and macro F1 score. The results are summarized in Table 3. The F1 score is a metric that balances precision and recall by calculating their weighted average or harmonic mean. As the magnitudes of macro precision and recall in each model are similar to the corresponding macro F1 score, only accuracy and macro F1 score are presented and discussed hereafter.

**Table 3:** Prediction performance of machine learning and deep learning models under hold-out and leave-cow-out cross-validation (CV) designs. KNN, RF, GB, MLP, and CNN refer to k-nearest neighbor, random forest, gradient boosting classifier, multi-layer perceptron, and convolutional neural network, respectively.

| CV designs | Scenarios | Models | Accuracy | Precision | Recall | F1 score |
| --- | --- | --- | --- | --- | --- | --- |
| **Hold-out** | 9:1 | **KNN** | 0.95 | 0.94 | 0.95 | 0.95 |
|  |  | **GB** | 0.96 | 0.95 | 0.95 | 0.95 |
|  |  | **RF** | 0.95 | 0.93 | 0.94 | 0.94 |
|  |  | **MLP** | 0.91 | 0.91 | 0.9 | 0.9 |
|  |  | **CNN** | 0.91 | 0.9 | 0.9 | 0.9 |
|  | 8:2 | **KNN** | 0.95 | 0.94 | 0.95 | 0.94 |
|  |  | **GB** | 0.95 | 0.94 | 0.95 | 0.94 |
|  |  | **RF** | 0.94 | 0.93 | 0.94 | 0.93 |
|  |  | **MLP** | 0.91 | 0.9 | 0.9 | 0.9 |
|  |  | **CNN** | 0.92 | 0.91 | 0.91 | 0.91 |
|  | 7:3 | **KNN** | 0.95 | 0.94 | 0.94 | 0.94 |
|  |  | **GB** | 0.95 | 0.94 | 0.95 | 0.94 |
|  |  | **RF** | 0.94 | 0.93 | 0.94 | 0.93 |
|  |  | **MLP** | 0.91 | 0.9 | 0.92 | 0.91 |
|  |  | **CNN** | 0.92 | 0.9 | 0.91 | 0.9 |
|  | 6:4 | **KNN** | 0.94 | 0.93 | 0.94 | 0.94 |
|  |  | **GB** | 0.95 | 0.93 | 0.95 | 0.94 |
|  |  | **RF** | 0.94 | 0.92 | 0.94 | 0.93 |
|  |  | **MLP** | 0.91 | 0.89 | 0.91 | 0.9 |
|  |  | **CNN** | 0.91 | 0.9 | 0.91 | 0.9 |
|  | 5:5 | **KNN** | 0.94 | 0.93 | 0.94 | 0.93 |
|  |  | **GB** | 0.95 | 0.93 | 0.94 | 0.94 |
|  |  | **RF** | 0.94 | 0.92 | 0.94 | 0.93 |
|  |  | **MLP** | 0.9 | 0.89 | 0.9 | 0.89 |
|  |  | **CNN** | 0.91 | 0.9 | 0.89 | 0.9 |
| **Leave-cow-out** | Leave-one-cow-out | **KNN** | 0.79 | 0.68 | 0.66 | 0.65 |
|  |  | **GB** | 0.78 | 0.66 | 0.64 | 0.64 |
|  |  | **RF** | 0.82 | 0.73 | 0.72 | 0.7 |
|  |  | **MLP** | 0.79 | 0.67 | 0.65 | 0.64 |
|  |  | **CNN** | 0.81 | 0.68 | 0.67 | 0.66 |
|  | Leave-two-cow-out | **KNN** | 0.76 | 0.74 | 0.73 | 0.7 |
|  |  | **GB** | 0.73 | 0.78 | 0.7 | 0.82 |
|  |  | **RF** | 0.79 | 0.77 | 0.74 | 0.73 |
|  |  | **MLP** | 0.78 | 0.75 | 0.73 | 0.71 |
|  |  | **CNN** | 0.78 | 0.75 | 0.74 | 0.71 |
|  | Leave-three-cow-out | **KNN** | 0.72 | 0.73 | 0.7 | 0.67 |
|  |  | **GB** | 0.77 | 0.76 | 0.72 | 0.7 |
|  |  | **RF** | 0.77 | 0.77 | 0.73 | 0.71 |
|  |  | **MLP** | 0.82 | 0.81 | 0.78 | 0.76 |
|  |  | **CNN** | 0.76 | 0.74 | 0.73 | 0.7 |

In the hold-out CV design, five scenarios were evaluated by varying the training set size from 90% to 50%, where the overall performance ranged from 0.9 to 0.95 for accuracy and from 0.89 to 0.95 for F1 score. The best performance, in terms of both accuracy and F1 score, was achieved by the ML model GB, with an accuracy of 0.96 and an F1 score of 0.95 in the 90% training scenario. This was closely followed by two other ML models, KNN and RF, both with an accuracy of 0.95 and F1 scores of 0.95 and 0.94, respectively. The two DL models, MLP and CNN, both obtained an accuracy of 0.91 and an F1 score of 0.9. As the training set size decreased, both accuracy and F1 score slightly decreased, with ML models consistently showing slightly better performance than DL models.

In the LCO CV design, three scenarios were evaluated by leaving one, two, and three cows out of the training set, respectively. The prediction accuracy and F1 score ranged from 0.72 to 0.82 and from 0.64 to 0.82, respectively. In the L1CO scenario, the RF model returned both the highest accuracy of 0.82 and the highest F1 score of 0.7, followed by CNN with an accuracy of 0.81 and an F1 score of 0.66. In the L2CO scenario, RF also performed the best with an accuracy of 0.79, while MLP and CNN both attained an accuracy of 0.78. The highest F1 score in this scenario was obtained by GB (0.82), followed by RF (0.73). In the L3CO CV, MLP achieved the highest accuracy of 0.82 and F1 score of 0.76, followed by RF with an accuracy of 0.77 and an F1 score of 0.71. Generally, as the number of unseen cows in the testing set increased, accuracy tended to decrease, but there was no clear trend for the F1 score.

Overall, ML models slightly outperformed DL models across all scenarios in the hold-out CV design. However, this pattern did not hold consistently in the LCO CV design, where DL models, particularly MLP, showed comparable or even slightly better performance than ML models, especially when more unseen cows were present in the testing set. Overall performances of all models were higher in the hold-out CV design compared to the LCO CV design.

## 5 Discussion

This study explored the effect of different CV designs on the performance of ML and DL models in predicting cattle behavior from accelerometer data. Specifically, we applied three ML models and two DL models to predict multi-class cattle behavior using both hold-out and LCO CV designs. The main contributions of this study were to evaluate the impact of hold-out versus LCO CV on multi-class imbalanced cattle behavior prediction and to compare the prediction performance of ML and DL models.

### 5.1 Comparison of prediction performance between ML and DL models

This study applied three ML models, including KNN, GB, and RF, to evaluate the behavior prediction performance. In the hold-out CV, the three ML models achieved accuracies and F1 scores ranging from 0.94 to 0.96 and 0.93 to 0.95, respectively. These results were similar to or better than those reported in previous studies using ML models and hold-out CV designs [10, 11, 12, 15, 16, 32, 33]. For instance, Robert et al. [10] reported classification accuracies of 0.99, 0.98, and 0.68 for lying, standing, and walking behaviors, respectively, using logistic regression for 15 beef calves. Similarly, Dutta et al. [11] classified five major behaviors from 24 dairy cows using a bagging ensemble model and obtained an average accuracy of 0.96 and an F1 score of 0.89. Riaboff et al. [12] predicted four behaviors from 10 dairy cows using a decision tree model and reported an overall accuracy of 0.95 and an F1 score of 0.96. The two DL models in this study achieved accuracies and F1 scores both ranging from 0.9 to 0.91. Similarly, Balasso et al. [23] utilized a CNN model to classify five behaviors of 12 dairy cows and reported an overall accuracy and F1 score of 0.96. Liu et al. [34] developed a fully convolutional network for classifying seven behaviors from 12 beef cows, achieving an overall accuracy of 0.84. Our results showed that ML models performed consistently better than DL models. One potential reason could be that ML models are better suited for small datasets and simple patterns, whereas DL models typically require larger datasets to learn patterns and generalize well.

In the LCO CV, accuracies and F1 scores ranged from 0.72 to 0.82 and 0.64 to 0.82, respectively, for ML models, and from 0.76 to 0.82 and 0.64 to 0.76, respectively, for DL models. These accuracies and F1 scores aligned with recent studies. For example, Smith et al. [35] applied L3CO CV to classify grazing, ruminating, resting, and walking behaviors in 24 dairy cows and reported F1 scores of 0.98, 0.87, 0.85, and 0.73, respectively. Among the DL models in our study, the MLP model achieved better performance than the CNN model and showed comparable or better performance than ML models, particularly when more unseen cows were included in the testing set. This may be because the DL models can capture more complex patterns that encompass the diversity in behavior across different cows, allowing them to generalize better predictions for unseen cows than ML models. To the best of our knowledge, no previous studies have compared ML models and CNN models using LCO CV to predict cattle behavior, which prevents the direct comparisons of our results with earlier work. However, a recent study by Coelho Ribeiro et al. [19] applied RF and MLP models to predict the grazing activities of six Nellore bulls using L1CO CV and found an identical accuracy of 0.57 for both models. These findings are consistent with our results, where the MLP model showed similar performance comparable to ML models. This suggests that DL models, such as MLP, may be robust for predicting behavior in scenarios involving new animals, but further investigation with a more comprehensive dataset is needed.

### 5.2 Impact of CV designs on prediction performance

Our results indicated that the prediction performance of both ML and DL models is much lower when using LCO CV compared to hold-out CV. The hold-out CV design used in this study is a form of random CV that involves random partitioning of behavior sensor data into training and testing sets. This random partition can allow the behavior records from the same cow to be included in both sets, introducing dependence or covariance between the sets. Notably, such dependence or covariance is often referred to as data leakage in CV and has been reported in other fields [36, 37]. As a result, the prediction performance from the hold-out CV may be inflated or overoptimistic due to model overfitting. Roberts et al. [17] reviewed and discussed how random CV may lead to serious underestimation of predictive error and result in overly optimistic prediction when applied to ecological data with temporal, spatial, hierarchical, or phylogenetic structure if these structures are ignored. Using simulations and case studies, the authors demonstrated that block CV, which strategically partitions data rather than randomly, is generally more suitable for predicting new data. Similarly, Wang and Bovenhuis [18] investigated the use of milk infrared spectra data to predict methane emissions using both random and block CV designs. They found that random CV produced overly optimistic predictions compared to block CV using farms as blocks, likely due to the confounding effect of farms. Likewise, Coelho Ribeiro et al. [19] compared grazing activity prediction using wearable sensor data under random and block CV designs and emphasized the importance of using management knowledge to strategically split data, as the seemingly good predictions from random CV were likely due to data dependence. In the current study, we used the LCO CV to assign all behavior records from specific cows to the testing set and used records from other cows as the training set. The LCO CV used here is a form of block CV, where we strategically partitioned data using cows as blocks to ensure that predictions for the testing set were made on unseen cows, avoiding dependence or covariance from the same cow. This explains why our prediction performance from LCO CV is much lower than that from hold-out CV, which was also consistent with other recent studies that also reported lower performance from block CV compared to random CV [21, 38].

Accelerometer data have been widely used in previous studies to predict cattle behavior, with most applying random CV to evaluate the prediction performance. However, these data are usually recorded sequentially over time for each cattle, creating temporal dependencies within a cattle’s data. Randomly partitioning data as shown in our hold-out CV ignores these dependencies and can split data from the same cattle into both training and testing sets, potentially leading to overly optimistic prediction performance. Our comparisons of prediction performance between LCO and hold-out CV designs suggest that block CV may better account for dependencies within the data, making it a potentially more appropriate approach when such dependencies exist. This could be particularly useful for scenarios involving predictions for unseen animals or generalizing predictions from one farm to another.

## 6 Conclusion

The current study investigated how CV designs influence the performance of ML and DL models in predicting multi-class imbalanced cattle behavior from accelerometer data. We found that prediction performance with hold-out CV was consistently better than with LCO CV. This is largely because the animal-specific dependence introduced by hold-out CV data partition artificially inflated the results. Despite the lower prediction performance, LCO CV, a form of block CV, could provide a more realistic and appropriate evaluation design when data exhibit inherent structure. We believe block CV should be used over random CV for evaluating the behavior prediction performance, particularly when predicting the behavior of unseen animals or new farms. Additionally, our results showed that the DL model, specifically the MLP, demonstrated comparable or better performance than other ML models in the LCO CV setting. This suggests that the MLP may be robust in capturing and predicting behavior across different animals, but further investigation with a more comprehensive dataset is needed for confirmation.

## CRediT authorship contribution statement

**Jin Wang:** Methodology, Software, Formal analysis, Visualization, Writing – original draft, Writing – review & editing. **Ziwen Yu:** Methodology, Writing – review & editing. **Ricardo C. Chebel:** Methodology, Writing – review & editing. **Haipeng Yu:** Conceptualization, Methodology, Writing – original draft, Writing – review & editing, Supervision, Project administration, Funding acquisition.

## Acknowledgments

The authors thank Dr. Gota Morota for the insightful discussions on the cross-validation designs used in this paper.

## Data availability

The tri-axial accelerometer dataset used in this study is a public dataset and is available at Zenodo.

