## Supplementary Figures S1, S2, and S3 for "Impact of cross-validation designs on cattle behavior prediction using machine learning and deep learning models with tri-axial accelerometer data"

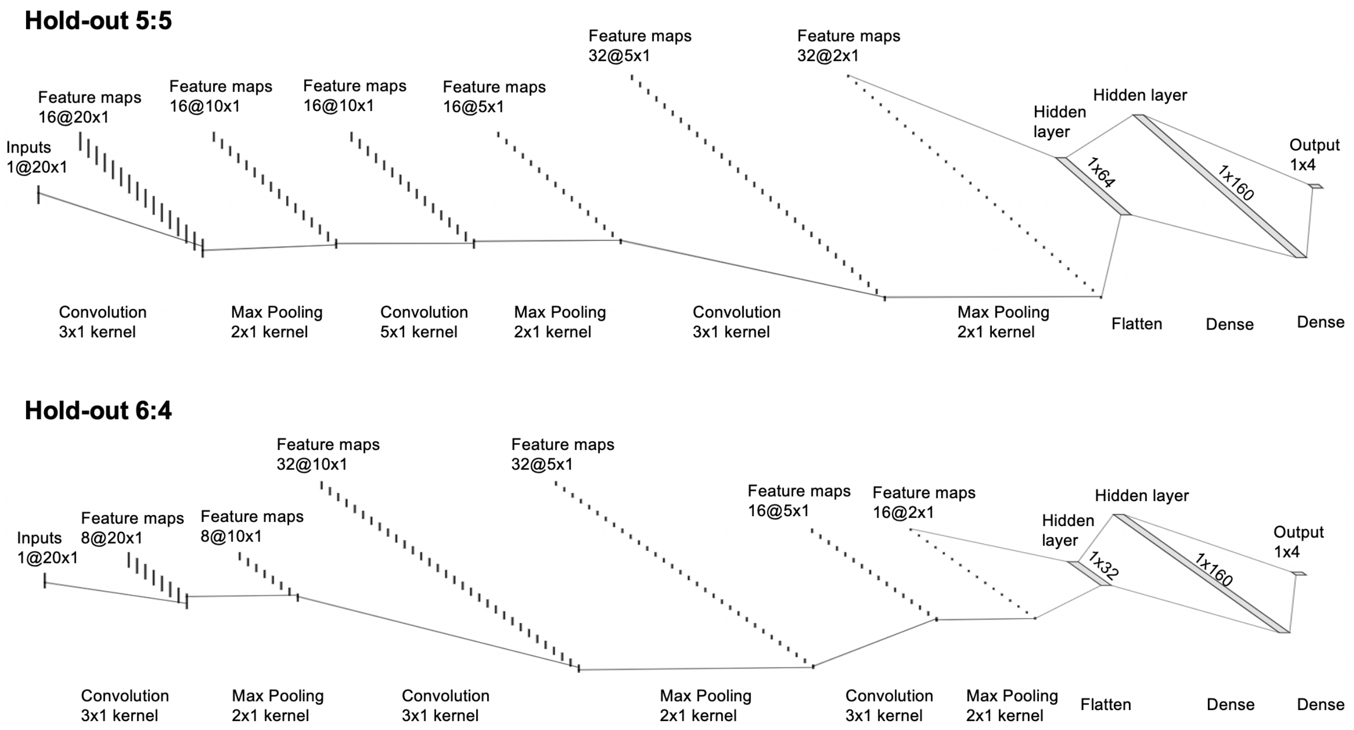


Figure S1: Convolutional neural network structures for hold-out (5 : 5) and hold-out (6 : 4) designs after hyperparameter tuning.

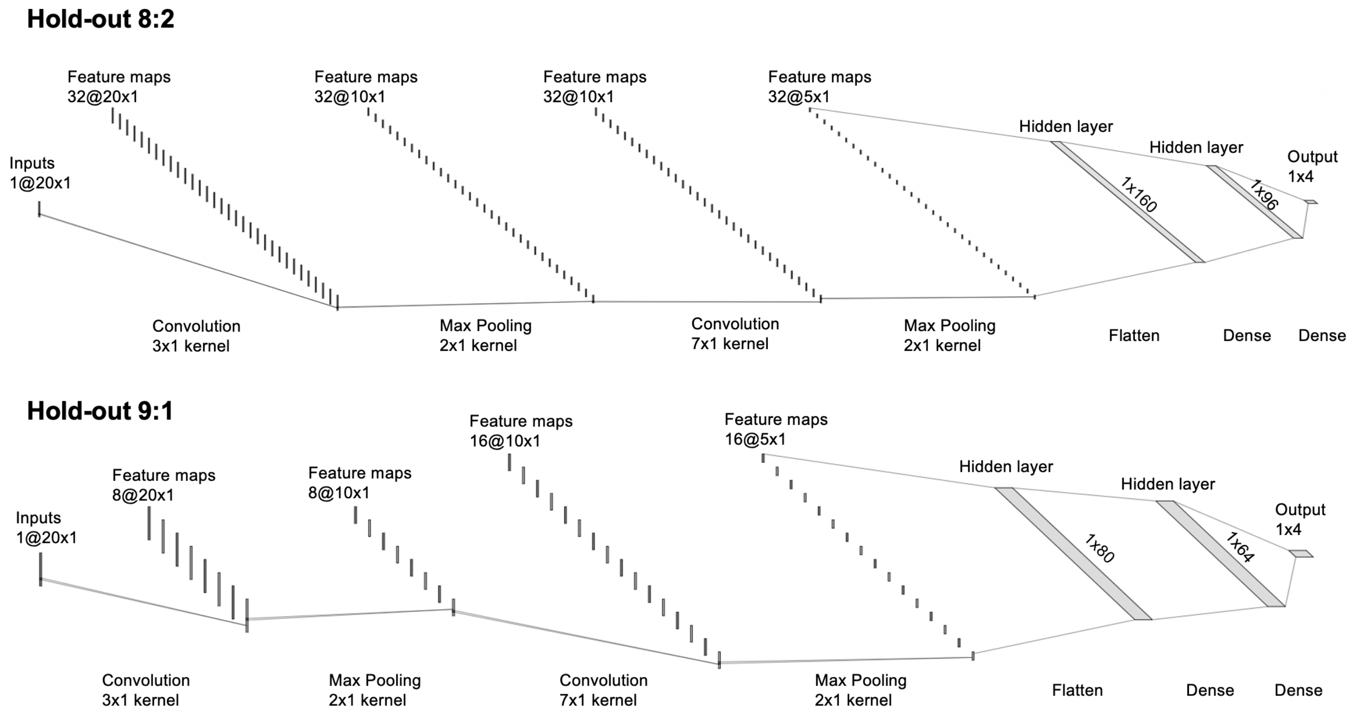


Figure S2: Convolutional neural network structures for hold-out (8 : 2) and hold-out (9 : 1) designs after hyperparameter tuning.

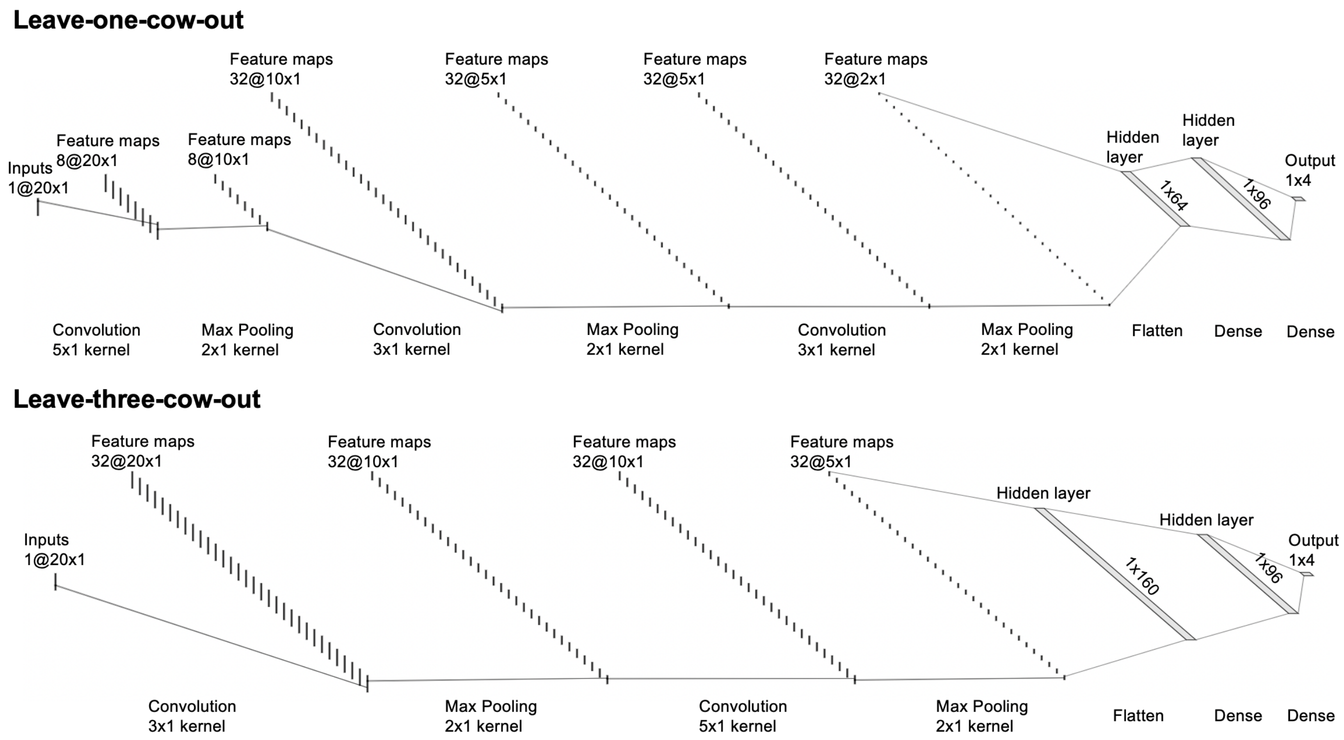


Figure S3: Convolutional neural network structures for leave-one-cow-out and leave-three-cow-out cross-validation designs after hyperparameter tuning.
